# GeneToList: A web application to assist with gene identifiers for the non-bioinformatics-savvy scientist

**DOI:** 10.1101/2022.06.09.494882

**Authors:** Joshua D. Breidenbach, E. Francis Begue, David J. Kennedy, Steven T. Haller

## Abstract

The increasing incorporation of omics technology into clinical and translational medicine presents challenges to end users of the large and complex datasets that are generated by these methods. A particular challenge in genomics is that the nomenclature for genes is not uniform between large genomic databases or between commonly used genetic analysis tools. Furthermore, outdated genomic nomenclature can still be found amongst scientific communications including peer-reviewed manuscripts. Therefore, a web application (GeneToList) was developed to assist is gene ID conversion and alias matching, with a specific focus on achieving a user-friendly interface for the non-bioinformatics-savvy scientist.

**Availability and implementation:** GeneToList is available at https://www.genetolist.com/. The tool is a web application that is compatible with many standard browsers.

## 1. Introduction

The increasing popularity of omics technologies in biomedical research has led to the birth of a subfield of data science, bioinformatics. While these techniques are becoming crucial to research, it is important to recognize that not all who stand to benefit from these advancements are poised to learn programming languages or become bioinformaticians. Additionally, the myriad of information attained through methods such as next-generation sequencing – based RNA-sequencing requires the community to constantly update gene and protein nomenclature. When dealing with the complex datasets generated by these methods and attempting to utilize the many genetic analysis tools available, there is difficulty in matching the format of one output to the required input of another. Additionally, obsolete genomic nomenclature persists colloquially and amongst peer-reviewed manuscripts. While great efforts have been made to allow for the conversion of gene identifiers, these usually require advanced knowledge of programming languages (biomaRt, MyGene - https://mygene.info/, and org.Hs.eg.db) [1, 2]. Otherwise, there are a few web applications which provide a user interface for the conversion of gene IDs. However, some are intended as an initial step of a more complex and powerful tool instead of a dedicated application for this purpose (DAVID - https://david.ncifcrf.gov/home.jsp) [3]. Others are dedicated, but rely on specific user input such as the input ID type and desired output, which may be a barrier for the unfamiliar scientist (g:Convert - https://biit.cs.ut.ee/gprofiler/ and bioDBnet - https://biodbnet-abcc.ncifcrf.gov/db/db2db.php) [4, 5]. Importantly, the authors are not aware of any tool which assists in alias matching especially in situations when obsolete IDs are ambiguous. Therefore, we set out to create a web application with a graphical user interface that can assist in the conversion of gene IDs and that disambiguates obsolete gene IDs in a high throughput manner suitable for large lists of genes.

## 2. Materials and Methods

### 2.1 Data Collection

Gene information for more than 34,000 taxa were collected from the National Center for Biotechnology and Information (NCBI) Gene resource [6-8]. Therefore, the application supports any taxa with gene information stored by NCBI, including archaea, fungi, invertebrates, mammalian and non-mammalian vertebrates, plants, protozoa, and viruses. Supported databases of gene IDs include NCBI Gene Symbols, NCBI Gene IDs (Entrez IDs), OMIM IDs, HGNC IDs, Ensembl IDs, and more taxa specific identifiers.

### 2.2 Application

This web application assists in 2 separate tasks. The first, is disambiguating obsolete gene nomenclature. A single search term or a list of terms (separated by comma or white space) can be entered into the text box and added to an existing list, or used to begin a new one. Searched terms are first matched as-is with a database of gene information for the selected taxonomy. Exact matched as added directly to the Final List. Additionally, matches after only slight alterations such as case changes, hyphenation, or removal of Greek letters are marked as “Auto-accepted Suggestion” and added to the Final List. More ambiguous terms are compared with gene synonyms and those with any potential matches are marked in the Final List and await the user to make a decision. Searched terms with ambiguous matches are selected one-at-a-time from a dropdown and their suggestions are listed along with other gene synonyms and descriptions. The most likely suggestion (based on the exact match of the searched term with a synonym) will be listed first. Lastly, those without any matches or those which are duplicate terms are marked as “No Match”, or “Duplicate Term” in the Final List, respectively. This functionality assists in the curation of a list of genes with officially recognized uniform identifiers.

The second task that the application assists with is the conversion of gene identifiers between formats, such as Ensembl ID’s and official gene symbols recognized by NCBI. Simply by entering a gene or list of genes into the input field, a list of curated genes is returned to the user as a table. In this way, gene IDs are converted without requiring the user to select the input type.

There are options to adjust the information included in the Final Table and the user can save it as a CSV file. Additionally, there are options to directly copy the matched NCBI gene symbols, NCBI gene ID (Entrez), or the full table to the clipboard. Users may add genes to their curated Final List through multiple iterations of searches. Importantly, the total input and output lists will be the same order and length, to eliminate confusion in the case of large input lists. Finally, the application provides links to follow-up analysis such as ontology (PantherDB.org) that may be of interest now that the user has a curated and/or converted list of uniform gene IDs.

### 2.3 Implementation

GeneToList was built as a web application in Python (3.8) using the Plotly Dash package (2.0.0), which provides a Python framework for web applications and relies on common javascript web frameworks Flask (2.0.2), Plotly.js (5.5.0), and React.js. GeneToList is compatible with many modern browsers on desktop and mobile including Google Chrome, Mozilla Firefox, Microsoft Edge, and Safari.

## 3. Results

### 3.1 Gene Alias Disambiguation

To demonstrate GeneToList’s capacity for disambiguation of obsolete or otherwise unofficial gene identifiers, we investigated a list of common inflammation related genes retrieved from a recent publication [9]. These 10 genes were searched in GeneToList by the names by which they were referred (see “Searched Terms” in Table 1). GeneToList found 4 exact matches, 1 automatically accepted suggestion, and 5 suggestions which required the user’s decision. For example, *IL-8* had the suggestions: *CXCL8, CXCR1, CXCR2*, and *CXCR2P1*. Upon evaluation of the common synonyms listed, we determined that *CXCL8* was the best match. Additionally, because the algorithm in GeneToList found *CXCL8* to be the most likely choice, it was listed as the top suggestion. After similar suggestion selection for the remaining searched terms, we were left with a list of matched symbols (see “Matched Symbol” in Table 1).

**Table 1.** Results of an example search of pro-inflammatory genes with GeneToList, demonstrating the disambiguation of gene IDs.

| Searched Term | Match Type | Matched Symbol |
| --- | --- | --- |
| TGF-β | Suggestion Accepted | TGFB1 |
| IL-8 | Suggestion Accepted | CXCL8 |
| MCP-1 | Suggestion Accepted | CCL2 |
| CRP | Exact Match | CRP |
| TNF-α | Suggestion Accepted | TNF |
| CXCR1 | Exact Match | CXCR1 |
| CXCR2 | Exact Match | CXCR2 |
| CCR2 | Exact Match | CCR2 |
| MYPT1 | Suggestion Accepted | PPP1R12A |
| TGF-β1 | Auto-accepted Suggestion | TGFB1 |

### 3.2 Gene ID Conversion

Because of the disambiguation feature of GeneToList, it is able to serve the purpose of a gene ID converter with better outcomes than other common ID converters. To demonstrate this, we used the same list of Searched Terms as in Table 1 and attempted conversion to Entrez IDs in GeneToList, g:Convert, DAVID and bioDBnet. These results are summarized in Table 2.

**Table 2.** Results of gene ID conversion from GeneToList and other common conversion tools.

| Searched Term | GeneToList | g:Convert | DAVID | bioDBnet |
| --- | --- | --- | --- | --- |
| TGF-β | 7040 | - | - | - |
| IL-8 | 3576 | - | - | - |
| MCP-1 | 6347 | - | - | - |
| CRP | 1401 | 1401 | 1401 | 1401 |
| TNF-α | 7124 | - | - | - |
| CXCR1 | 3577 | 3577 | 3577 | 3577 |
| CXCR2 | 3579 | 3579 | 3579 | 3579 |
| CCR2 | 729230 | 729230 | 729230 | 729230 |
| MYPT1 | 4659 | - | - | - |
| TGF-β1 | 7040 | - | - | - |

While GeneToList returned Entrez IDs for all searched terms, the other tools were only able to return 4 out of 10 queried. It is important to note that when GeneToList was used first to disambiguate the terms (such as in Table 1), and then the “Matched Symbol” list was run through these other conversion tools, Entrez IDs were found for all (Not Shown). This is an example of the utility of disambiguation before follow-up workflow.

## 4. Conclusion

The result of these efforts is a publicly available and free to use web application (GeneToList; https://www.genetolist.com/) to assist biologists and biomedical scientists in navigating gene data. This tool assists in disambiguation of gene IDs and was found to yield better results in ID conversion compared with other common gene ID conversion tools. This is meant to aid in the uniformity of a list of genes before being used for any following analysis.

